# AlignPCA-2D: PCA-Reduced Euclidean Vector Alignment for 2D Classification in Cryo-EM

**DOI:** 10.64898/2026.02.11.705027

**Authors:** E. Ramírez-Aportela, O. L. Zarrabeitia, Y. C. Fonseca, T. Ceska, S. Subramaniam, J.M. Carazo, C.O.S. Sorzano

**Affiliations:** Centro Nacional de Biotecnología (CSIC), C/ Darwin, 3, 28049, Cantoblanco, Madrid, Spain; Gandeeva Therapeutics, Inc., Vancouver, BC, Canada; Department of Biochemistry and Molecular Biology, University of British Columbia, Vancouver, BC, Canada

## Abstract

Cryogenic Electron Microscopy (cryo-EM) has transformed structural biology by enabling high-resolution reconstruction of macromolecular complexes from noisy projection images. However, the intrinsic heterogeneity and low signal-to-noise ratio of cryo-EM datasets make 2D classification a critical and computationally demanding step in the processing workflow. Here, we introduce AlignPCA-2D, a PCA-space Euclidean vector alignment method for fast, interpretable 2D classification in cryo-EM. By projecting particle images and class representations into a compressed latent PCA space, AlignPCA-2D reduces data dimensionality while preserving meaningful structural variability. The image-to-class assignment is then performed using Euclidean distance, enabling efficient and accurate classification. We benchmark AlignPCA-2D against established cryo-EM software, such as RELION and cryoSPARC, and demonstrate that it achieves competitive alignment accuracy while substantially reducing computational cost. This approach provides a lightweight alternative for large-scale 2D classification tasks, and its modular design makes it compatible with existing cryo-EM processing pipelines.

## Introduction

Cryogenic Electron Microscopy (cryo-EM) has emerged as a transformative technique for resolving the structures of biological macromolecules in near-native conditions. By capturing two-dimensional projections of molecules frozen in vitreous ice, cryo-EM provides insights into structural dynamics that are often inaccessible by other methods. However, raw cryo-EM datasets typically contain heterogeneous mixtures of particle conformations and orientations, along with significant levels of noise, making the extraction of meaningful structural information a considerable computational challenge. Two-dimensional (2D) classification plays a pivotal role in cryo-EM image processing pipelines. It groups similar particle images based on shared structural features, enhancing signal-to-noise ratio and revealing distinct molecular states and orientations. This step improves data interpretability and serves as a crucial foundation for subsequent processes, such as generating ab initio 3D models and refining the volume.

A central difficulty in 2D classification arises from the high dimensionality of cryo-EM images, which can contain hundreds of thousands of pixels per particle. Analyzing these data directly is computationally intensive and susceptible to overfitting. To mitigate these challenges, dimensionality reduction techniques—particularly Principal Component Analysis (PCA)—have been widely adopted. PCA projects high-dimensional image data onto a lower-dimensional subspace defined by the principal components, capturing the dominant modes of variation while suppressing noise and redundancy. This approach accelerates computation and also enhances the interpretability of structural variability within the dataset. To quantify structural similarity among images, different metrics have been explored, including cross-correlation and Euclidean distance. While cross-correlation has traditionally been employed in cryo-EM alignment, simpler geometric metrics in reduced latent spaces offer promising computational advantages.

In this work, we introduce AlignPCA-2D, a novel method for 2D classification in cryo-EM that combines PCA-based vector compression with Euclidean distance for image-to-class assignment. By operating directly in a compressed latent space, AlignPCA-2D achieves fast, interpretable, and accurate classification without sacrificing structural fidelity. We evaluate its performance across multiple publicly available datasets and benchmark it against two leading cryo-EM tools, RELION (Scheres, 2012) and CryoSPARC (Punjani et al., 2017), demonstrating comparable or superior performance with substantially reduced computational cost.

## 2 Methods

The proposed AlignPCA-2D algorithm is inspired by the classical *k* -means clustering paradigm, whose essence lies in a two-step iterative procedure. In the first step, each particle image is compared with a set of current class representatives (or prototypes) and assigned to its most similar one according to a chosen distance metric—a process referred to here as *2D image alignment*. In the second step, the class representatives are updated by averaging all particle images assigned to each class, thus producing new prototypes that better reflect the underlying structural variability. These two stages—alignment and representative update—are iterated until convergence, progressively improving both the class assignments and the accuracy of the class averages.

### 2.1 2D Image alignment

It is very common to quantify the difference between two images by a weighted quadratic distance in vector space. Let **x, y** ∈ ℂ^*n*^ represent two images written as column vectors, and let *W* be a positive definite weighting matrix. The *weighted squared distance* is then defined as

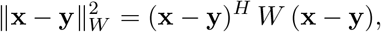

where the superscript *H* denotes the Hermitian transpose.

Let 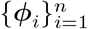 be an orthonormal basis of ℂ^*n*^ and let Φ = [ ***ϕ***_1_ … ***ϕ***_*n*_ ] be the unitary matrix whose columns are the basis vectors, so that Φ^*H*^ Φ = *I*. Any image **x** can be expanded as **x** = Φ**a**, where **a** is the vector of coefficients in this basis. Similarly, **y** = Φ**b**.

For a general Hermitian positive semidefinite weighting matrix *W*, the weighted quadratic distance between **x** and **y** is

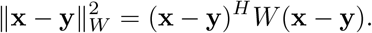

Substituting the expansions **x** = Φ**a** and **y** = Φ**b** gives

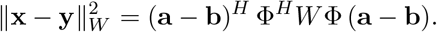

Therefore, the weighted distance can be equivalently expressed in the space of orthogonal coefficients as

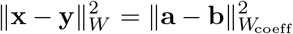

with *W*_coeff_ = Φ^*H*^ *W* Φ. This identity is the general expression underlying Parseval’s theorem: an orthogonal transformation preserves inner products and, consequently, weighted norms up to a change of metric. It holds for any orthogonal expansion, including (i) the real-space representation, where Φ is the identity matrix (the canonical basis of Kronecker’s deltas); (ii) the Fourier representation, where Φ is the unitary discrete Fourier transform matrix composed of complex exponentials; and (iii) the principal component representation, where Φ diagonalizes the covariance matrix of the data, providing an orthonormal basis that concentrates the variance in a few dominant modes. Hence, the same metric structure applies consistently across spatial, frequency, or statistical bases, ensuring that image distances can be computed equivalently in any of these domains.

If *W* happens to be diagonal in the chosen basis, then *W*_coeff_ = *W* = diag(*w*_*i*_) and the expression simplifies to

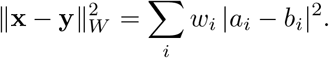

It is important to emphasize that there is no unique or “correct” choice for the weighting matrix *W* . Any selected form of *W* merely reflects our preferences regarding how to emphasize or de-emphasize particular components of the data—such as spatial regions, frequency bands, or principal components—during distance computation. Different weighting schemes encode different assumptions about which aspects of the images are considered more reliable or informative. Still, none can be regarded as intrinsically superior from a purely mathematical standpoint. In practice, the optimal choice of *W* depends on the specific goals of the analysis, the characteristics of the dataset, and the desired trade-off between robustness to noise and sensitivity to structural detail.

In practical applications using PCA, however, it is often convenient to restrict the representation to a limited number of coefficients. For instance, one may truncate the expansion to the first *K* components that capture most of the variance in the data:

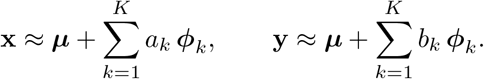

In this case, the distance becomes only an approximation to the full weighted distance, since the contributions from the discarded components are neglected. Formally, this truncation can be viewed as using a weighting matrix *W* with zeros along the directions corresponding to the omitted components, that is, a positive semidefinite (rather than positive definite) *W* . Consequently, the computed distance quantifies similarity only within the retained subspace, effectively filtering out the components associated with noise or minor variations.

In what follows, we assume that both images are in Fourier space. It is convenient to think that *W* is a diagonal matrix representing some preferences on the relative importance of the frequencies to compute the distance. We chose *W* to be a filter that gives more weight to high frequencies, so that they have a greater influence on the alignment, and we achieve more accuracy in the class representative. However, the input images have more noise at high frequencies, so we prefer slightly dampening those before doing the comparison, that is, we compare the images as

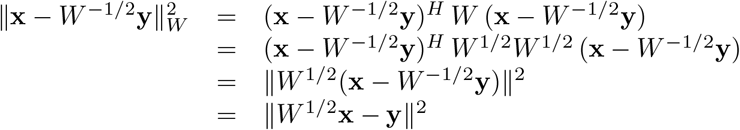

Additionally, as discussed above, computing the distance in the real, Fourier, or PCA domains is completely equivalent, since these are all orthogonal representations related by unitary transforms. In practice, we compute the distance in the PCA space of the Fourier components: two independent PCAs are performed, one for the real part and another for the imaginary part of the Fourier coefficients. The analysis is restricted to a specific frequency band, which is mathematically equivalent to setting the weights of the components outside that band to zero. This band limitation is determined empirically by evaluating the Fourier Ring Correlation (FRC) of the class, which provides an estimate of the spatial frequency range containing reliable structural information.

Finally, given an experimental image **y** and a set of reference images {**x**_*i*_}, the alignment problem consists in finding both the reference and the geometric transformation that best match **y**. Each reference **x**_*i*_ can be transformed according to a set of parameters ***θ*** (e.g., in-plane rotation and translation), yielding the transformed image **x**_*i*_(***θ***). The optimal alignment is then obtained by jointly minimizing the weighted distance over both the transformation parameters and the reference index:

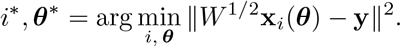

As an implementation note, we use the vectors *W* ^1*/*2^**x**_*i*_ as the class representatives, denoted **C**_*i*_. Then, the comparison above becomes

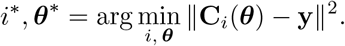

In the iterative process of assigning images to classes based on their distance and then updating the class representatives, we generate, at each iteration, an exhaustive grid of in-plane rotations and translations for every representative. Given the best alignment parameters from the previous iteration, the input images are pre-corrected accordingly, and the new in-plane parameters are combined with the previously found ones.

### 2.2 Representative updates

The refinement procedure proceeds in two stages. In the first stage, a subset of *N*_1_ selected particles are used to initialize and iteratively refine the class averages. Initial reference classes are obtained using *k* -means clustering applied to the PCA-compressed particle representations (with *k* equal to the number of requested classes). This unsupervised partitioning yields a set of structurally coherent class prototypes that serve as initial references for subsequent refinement aimed at improving intra-class consistency and overall representational accuracy. At each iteration, all particles are compared against the current set of class references, and each particle is assigned to its most similar class according to the distance in PCA-Fourier space. The particles assigned to the same class are then averaged, as in standard *k* -means, and the resulting averages are filtered by *W* ^1*/*2^ to generate the updated class references for the next iteration. Once the classes stabilize in the first stage, the remaining particles are processed in batches of size *N*_*b*_ and assigned to their most similar reference classes, again based on the distance in PCA-Fourier space. In the second stage, the class representatives obtained from stage 1 are further refined using an RMSProp-inspired optimization scheme. Let **C**_*t*_ denote the class representation at iteration *t*, and let **g**_*t*_ be the gradient of the cost function, which is defined as the sum of the distances between all images assigned to that class and the current class representation:

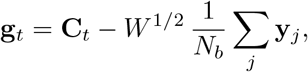

where the sum runs over the *N*_*b*_ particles in the current batch. The moving average of the squared gradient, **v**_*t*_, is then updated as

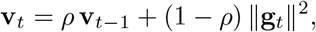

where *ρ* is the decay factor. The class representation is subsequently updated according to

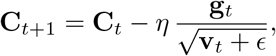

where *η* is the learning rate, *ϵ* is a small constant introduced for numerical stability, and the division between vectors is performed component-wise.

This iterative refinement strategy allows the class representations to evolve smoothly as new particle batches are incorporated, yielding stable and progressively more accurate class averages that capture the structural variability present in the dataset. The batch updates make our algorithm naturally suited for streaming workflows. As modern cryo-EM facilities achieve faster image acquisition and near real-time data transfer, the ability to process data in parallel with acquisition has become increasingly relevant. Real-time classification enables on-the-fly quality assessment, provides rapid feedback for microscope operators, and facilitates early detection of sample or alignment issues, thereby enhancing the overall efficiency of cryo-EM data collection.

## Code availability

AlignPCA-2D has been developed in Python using PyTorch libraries and is publicly available from Xmipp (de la Rosa-Trevín et al., 2013) in the development branches (which will eventually become the next release of Xmipp), and is integrated into the image-processing framework Scipion 3 (Conesa et al., 2023).

## 3 Results

To evaluate the performance of our method, we analyzed several publicly available cryo-EM datasets from EMPIAR. We compared the results with those obtained using two widely adopted tools, RELION and CryoSPARC. All datasets were processed within the Scipion 3 framework to ensure reproducibility and consistent data handling across methods.

We first imported the aligned micrographs into Scipion and estimated the CTF using both GCtf (Zhang, 2016) and CTFFind4 (Rohou and Grigorieff, 2015). A consensus of CTF parameters was then generated using Xmipp (Sorzano et al., 2021), retaining only micrographs with an estimated resolution better than 7°A. Particle picking was automatically performed with Cryolo (Wagner et al., 2019), followed by extraction and downstream 2D classification using the respective algorithms under comparison. For CryoSparc and RELION, default classification parameters were used to ensure a fair and reproducible comparison.

In our approach, CTF correction was applied using the Wiener filter implementation in Xmipp before PCA projection and classification. Only Fourier coefficients corresponding to spatial frequencies lower than 8°A were retained, as they contain the dominant alignment-relevant signal (Scheres, 2012). For all datasets, the number of PCA components retained in AlignPCA-2D corresponded to those that captured 75% of the total variance, ensuring a compact yet informative latent representation. Although this variance threshold was fixed, the actual number of components varied according to the intrinsic variability of each dataset. The extracted particle box size was set to 128 × 128 pixels in all cases to maintain consistency across datasets and methods.

### 3.1 Dataset Analysis

#### EMPIAR-10061

The first dataset, EMPIAR-10061 (Bartesaghi et al., 2015), contains raw cryo-EM images of *β*-galactosidase, a canonical benchmark in structural biology. A total of 1,539 micrographs were processed in Scipion (see Methods, Data processing section), yielding 381,792 automatically extracted particles used for 2D classification, with the number of classes fixed at 100 in all cases.

Figure 1 and Supplementary Figure 5 show the class averages obtained using AlignPCA-2D, whereas Supplementary Figure 1 shows a comparison of the classes obtained using CryoSPARC, RELION, and AlignPCA-2D. Additionally, Table 1 summarizes the computation times required for each method to generate 100 classes. When processing 381,792 particles, RELION requires more than four hours, whereas CryoSPARC and AlignPCA-2D completed the task in 30.11 minutes and 29.33 minutes, respectively.

**Figure 1.**
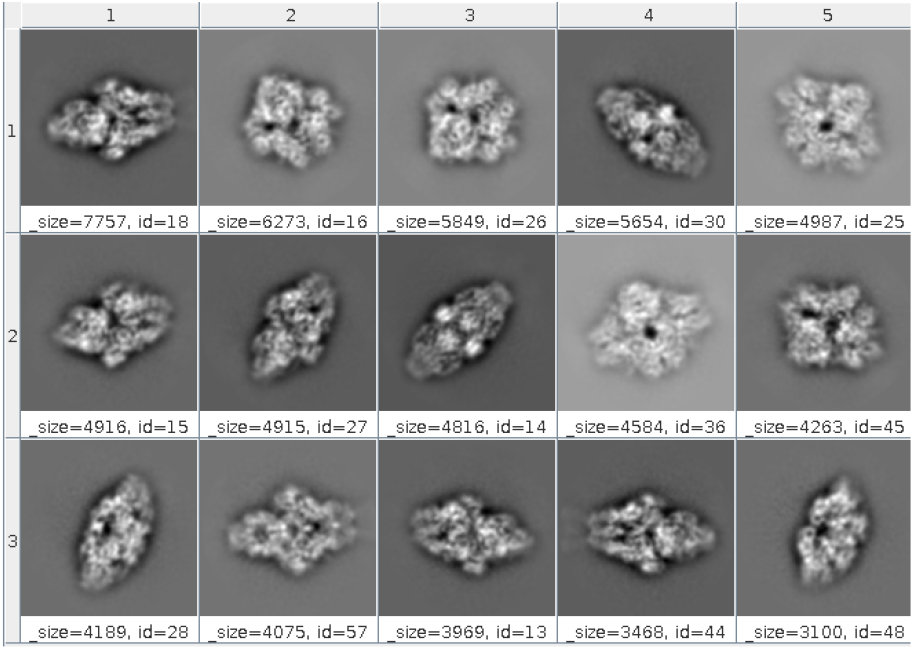
Representative 2D class averages from the EMPIAR-10061 dataset obtained with AlignPCA-2D.

**Table 1.**
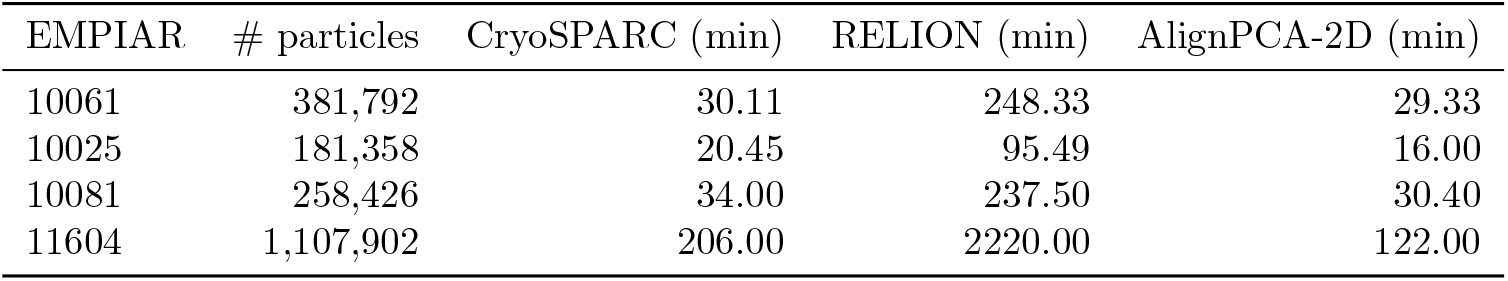
Comparison of computation times for different 2D classification methods.

| EMPIAR | # particles | CryoSPARC (min) | RELION (min) | AlignPCA-2D (min) |
| --- | --- | --- | --- | --- |
| 10061 | 381,792 | 30.11 | 248.33 | 29.33 |
| 10025 | 181,358 | 20.45 | 95.49 | 16.00 |
| 10081 | 258,426 | 34.00 | 237.50 | 30.40 |
| 11604 | 1,107,902 | 206.00 | 2220.00 | 122.00 |

Supplementary Figure 1 illustrates that all three methods successfully separated distinct particle orientations, while Supplementary Figure 5 demonstrates that AlignPCA-2D effectively discriminated signal from noise. Table 2 summarizes the number of particles retained after classification and the overlap between methods. Manual class selection was applied consistently across all approaches, retaining only classes that exhibited well-defined structural features and minimal noise. A total of 119,435 particles were selected by RELION, 132,408 by CryoSPARC, and 132,923 by AlignPCA-2D. Overlap between particle sets from different methods was computed using the consensus tools available in Scipion (Sorzano et al., 2021). The highest overlap was observed between CryoSPARC and AlignPCA-2D, with 80.8% of the total particles (132,923). The consensus between RELION and CryoSPARC reached 78.6%, while the overlap between RELION and AlignPCA-2D was 73.4%.

**Table 2.** Particle retention and overlap among 2D classification methods.

| EMPIAR | RELION | CS | PCA | CS-RELION (%) | CS-PCA (%) | RELION-PCA (%) |
| --- | --- | --- | --- | --- | --- | --- |
| 10061 | 119,435 | 132,408 | 132,923 | 104,050 (78.6) | **108,149 (82.7)** | 97,618 (73.4) |
| 10025 | 136,773 | 144,809 | 144,891 | **130,152 (89.9)** | **130,232 (89.9)** | 125,967 (86.9) |
| 10081 | 123,494 | 125,921 | 127,963 | 102,747 (81.6) | **108,497 (84.8)** | 100,913 (78.9) |
| 11604 | 405,267 | 332,065 | 332,679 | 235,419 (58.1) | **209,545 (63.0)** | 213,144 (52.6) |

So far, our analysis has focused on manually selected subsets of well-defined classes, which leave little room for ambiguity. Since this represents an early stage of the classification pipeline—where a relatively permissive selection is typically applied before a more stringent second round—it is expected that some classes may contain a small proportion of noisy or poorly aligned particles.

To further evaluate the particle-selection behavior of our method, we analyzed a subset of CryoSPARC classes (68 classes) that lacked well-defined structural features (i.e., “noise classes”), comprising 212,838 particles (Supplementary Figure 8). This set of particles was reprocessed using CryoSPARC, RE-LION, and AlignPCA-2D, and the resulting class averages are shown in Supplementary Figures 9, 10, and 11, respectively. As observed, neither CryoSPARC nor RELION produced any classes with well-defined structural features. In contrast, AlignPCA-2D successfully generated coherent classes containing approximately 23,707 well-aligned particles, corresponding to about 11,14% of the total dataset.

To assess the robustness of the alignments determined by AlignPCA-2D, we applied the orientations obtained by our method to the same particle subset. We re-ran both CryoSPARC and RELION using these alignment parameters as initial estimates. Under these conditions, CryoSPARC successfully aligned approximately 37,800 particles, producing interpretable class averages (Supplementary Figure 12), while RELION generated high-quality classes comprising roughly 39,300 particles (Supplementary Figure 13). A detailed inspection revealed that the majority of the particles recovered under these conditions were significantly displaced from the image center.

These results demonstrate that, despite comparable overall computation times, AlignPCA-2D provides a more exhaustive exploration of the rotational and translational spaces, enabling the recovery of particles that would otherwise be discarded or misaligned by CryoSPARC. Moreover, the reduced dependence of AlignPCA-2D on the initial centering accuracy of particle extractions makes it less sensitive to imperfections introduced during particle picking. Consequently, smaller extraction boxes can be employed without compromising classification quality, thereby reducing computational cost.

Importantly, by re-centering particles based on the alignment parameters obtained with AlignPCA-2D, particles initially rejected by other methods can be effectively reintegrated into downstream analyses. Together, these findings highlight that AlignPCA-2D enhances classification completeness and particle retention, providing a computationally efficient and robust alternative for the early stages of cryo-EM data processing.

#### EMPIAR-10025

The second dataset tested, EMPIAR-10025 (Campbell et al., 2015), consists of cryo-EM images of the T20S proteasome. A total of 195 micrographs were processed in Scipion, yielding 161,358 extracted particles. Figure 2 and Supplementary Figure 2 illustrate representative classes obtained using the different methods, while Supplementary Figure 6 shows the 100 classes obtained with AlignPCA-2D. The computation times, shown in Table 1, were comparable for CryoSPARC and AlignPCA-2D (17.48 minutes and 14.47 minutes, respectively), whereas RELION was significantly slower, requiring 95.49 minutes.

**Figure 2.**
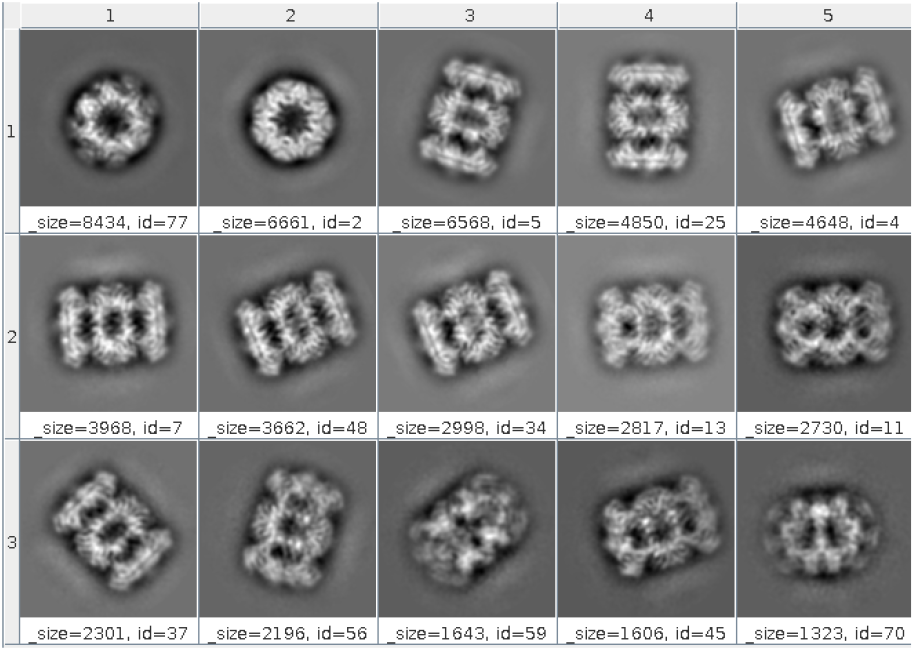
Representative 2D class averages from the EMPIAR-10025 dataset obtained with AlignPCA-2D.

The number of particles selected after classification was 136,773 for RE-LION, 144,809 for CryoSPARC, and 144,891 for AlignPCA-2D. Table 2 summarizes these results, along with the method overlap, which was close to 90% in all cases. On this dataset, AlignPCA-2D matches CryoSPARC in both iclassification quality and particle selection consistency, while maintaining a faster—or at least comparable— computational performance.

#### EMPIAR-10081 — HCN1 ion channel

The third dataset, EMPIAR-10081 (Lee and MacKinnon, 2017), corresponds to a membrane protein (HCN1 ion channel), which typically poses greater challenges due to structural flexibility and heterogeneity. After preprocessing, 258,426 particles were extracted.

The 2D classification performed with RELION required nearly four hours, whereas CryoSPARC and AlignPCA-2D completed the task in 34 minutes and 30 minutes, respectively. All methods successfully distinguished different particle orientations and produced high-quality class averages (Figure 3 and Supplementary Figure 3). The consensus analysis revealed a high degree of particle consistency, with an overlap of approximately 80% among the three methods (Table 2).

**Figure 3.**
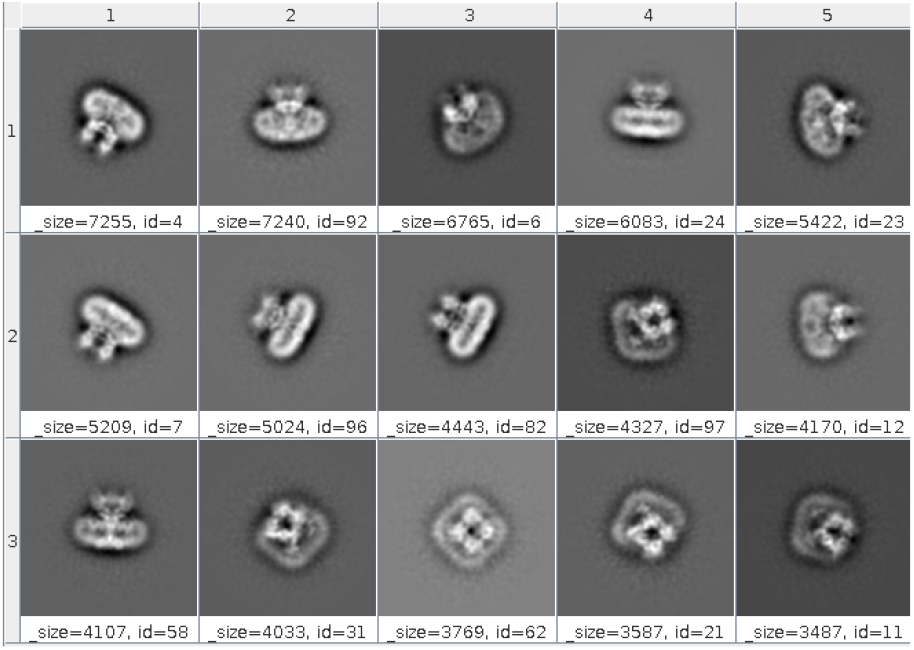
Representative 2D class averages from the EMPIAR-10081 dataset obtained with AlignPCA-2D.

Even for this high-complexity membrane protein dataset, AlignPCA-2D demonstrated robust classification performance (Supplementary Figure 7), generating well-defined class averages while maintaining computation times comparable to CryoSPARC and significantly faster than RELION.

#### EMPIAR-11604

The fourth dataset corresponds to the AP2 clathrin adaptor complex in the presence of heparin (Partlow et al., 2022). A total of 3,046 cryo-EM micrographs were preprocessed, resulting in the extraction of 1,107,902 particles for subsequent classification. This large-scale dataset is particularly well-suited for evaluating the scalability of AlignPCA-2D. For this dataset, 2D classification was performed with 400 2D classes.

Figure 4 and Supplementary Figure 4 show representative class averages obtained with the different methods, in which distinct particle orientations are clearly observed in all cases. Table 1 summarizes the computation times: CryoSPARC required 206 minutes (~ 3.4 h), and RELION required 2,220 minutes (37 h), whereas AlignPCA-2D completed the classification in only 122 minutes (∼2 h). This represents a speedup of approximately 1.7 × over CryoSPARC and 18 over RELION.

**Figure 4.**
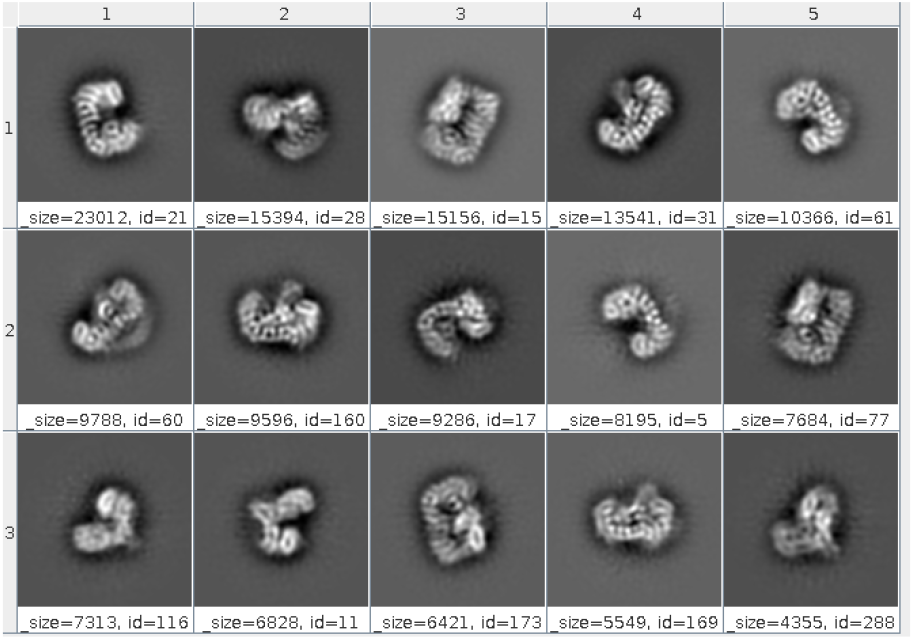
Representative 2D class averages from the EMPIAR-11604 dataset obtained with AlignPCA-2D.

Regarding particle consistency, Table 2 shows the overlap between the methods using our consensus approach. In this analysis, the selected particles exhibited moderate overlap of approximately 60%. This moderate overlap stems from the distinct underlying methods and search strategies utilized across the platforms. Despite these variations in particle selection, the class averages remain consistent, highlighting the complementary nature of these algorithmic approaches in capturing the dataset’s structural diversity. Overall, these results demonstrate that AlignPCA-2D achieves robust classification performance while maintaining excellent scalability, enabling efficient processing of large-scale cryo-EM datasets.

In the presented cases, we observed that all three tested methods successfully separated particles into distinct classes corresponding to different macromolecular orientations (Supplementary Figures 1–4). This demonstrates the robustness of our PCA-based algorithm for 2D classification. Additionally, our processing times were comparable to those of CryoSPARC and significantly faster than those of RELION. The highly competitive computation times achieved further highlight the efficiency of our method without compromising classification accuracy. Such efficiency is crucial for high-throughput cryo-EM workflows, where time savings can greatly enhance overall processing pipelines.

### 3.2 AlignPCA-2D in Streaming Cryo-EM Workflows

Traditional 2D classification in cryo-EM is typically performed as a static process, initiated once all particles have been collected. However, modern high-throughput data acquisition makes it increasingly desirable to perform classification in a streaming manner, allowing for real-time assessment of data quality and sample heterogeneity. To support real-time data analysis during cryo-EM acquisition, AlignPCA-2D was integrated into the Scipion framework for fully streaming operation. In this mode, an initial subset of particles is used to generate PCA-based reference classes. As new particles are acquired, they are automatically projected into the pre-learned PCA space and assigned to their most similar classes using Euclidean distance. Class averages are continuously refined through batch updates following the RMSProp scheme.

This implementation enables near–real–time monitoring of sample quality and orientation diversity, allowing users to detect issues such as mispicking or preferred orientations during acquisition.

## 4 Discussion

Dimensionality reduction has long been used in cryo-EM as a strategy to manage the high variability and large volume of particle datasets. In this work, we demonstrate that Principal Component Analysis (PCA) can be used not only as a data compression technique but also as an effective method for aligning and classifying data. By representing both particle images and class averages in a shared PCA space and assigning images through a Euclidean distance criterion, AlignPCA-2D achieves high-quality 2D classification with substantially reduced computational cost.

Results across multiple EMPIAR datasets show that AlignPCA-2D performs comparably to state-of-the-art tools such as RELION and CryoSPARC, while offering faster execution and a fully open-source implementation. Beyond its computational efficiency, AlignPCA-2D introduces a conceptual shift: alignment and classification in a reduced, well-defined latent space can be sufficient to capture the essential structural variability of cryo-EM data.

Its integration within Scipion and compatibility with streaming acquisition further extend its applicability, enabling real-time classification and adaptive feedback during data collection. The method’s open availability promotes transparency, reproducibility, and community-driven development—critical factors for advancing high-throughput cryo-EM workflows. Together, these properties position AlignPCA-2D as a lightweight, transparent, and accessible alternative for 2D classification, combining speed, interpretability, and open science.

