## Supplementary Figures for "AlignPCA-2D: PCA-Reduced Euclidean Vector Alignment for 2D Classification in Cryo-EM"

### Supplementary Figure 1

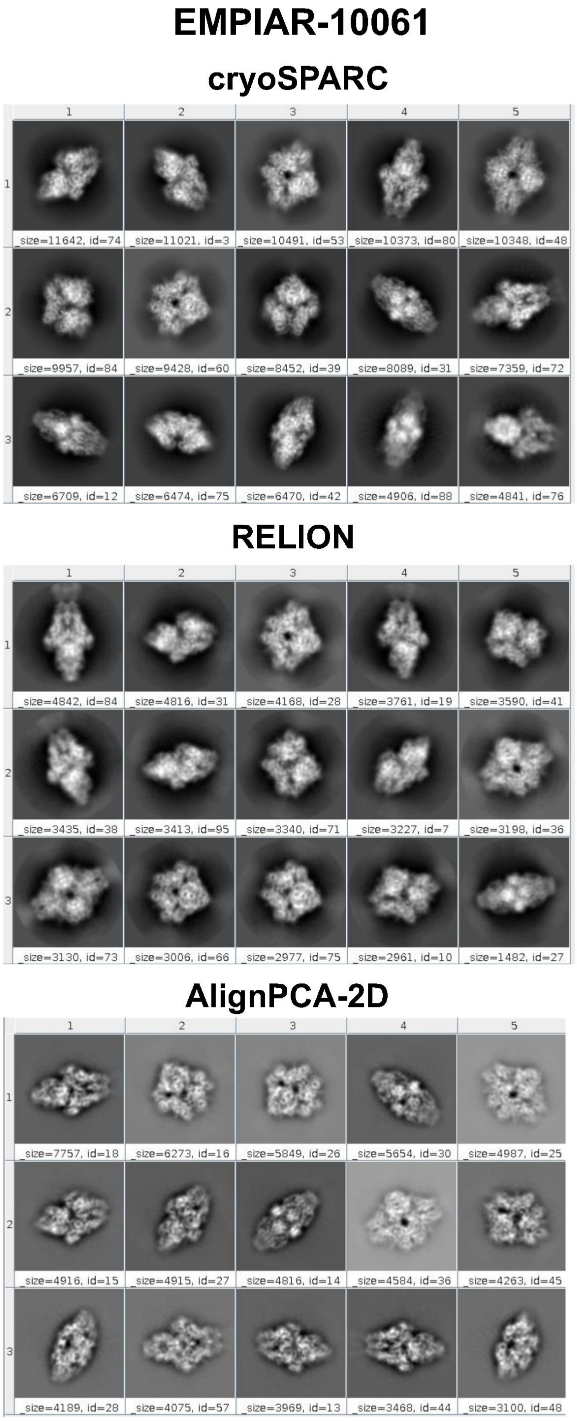


Supplementary Figure 1: Comparison of representative 2D class averages from the EMPIAR-10061 dataset obtained using CryoSPARC, RELION and AlignPCA-2D.

### Supplementary Figure 2

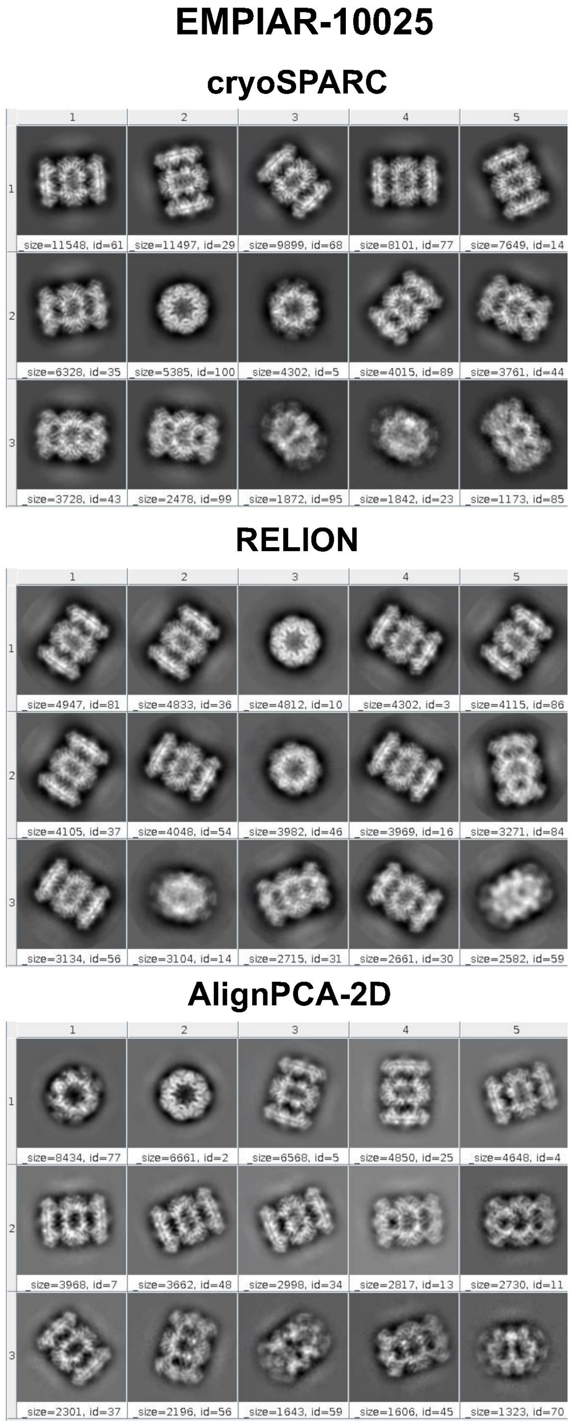


Supplementary Figure 2: Comparison of representative 2D class averages from the EMPIAR-10025 dataset obtained using CryoSPARC, RELION and AlignPCA-2D.

### Supplementary Figure 3

**EMPIAR-10081**

**cryoSPARC**

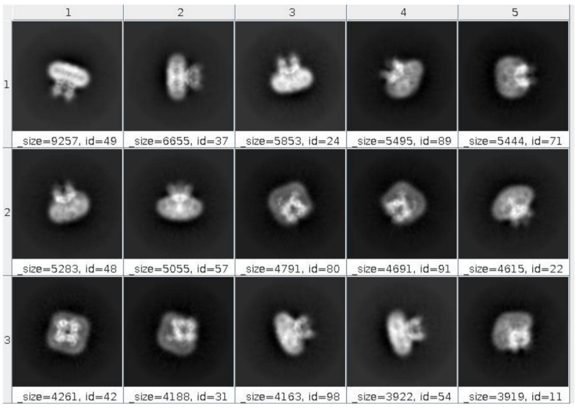


**RELION**

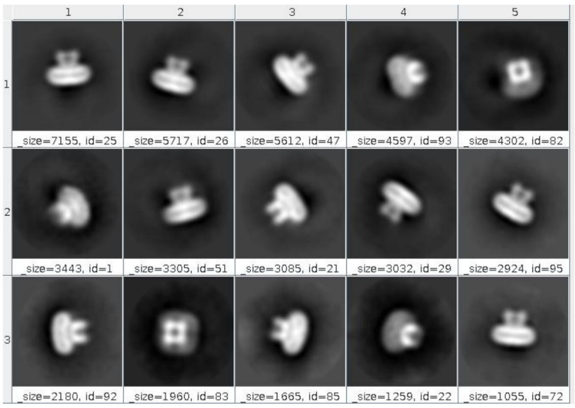


**AlignPCA-2D**

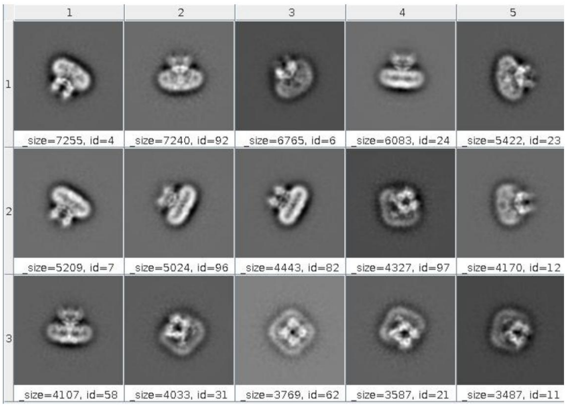


Supplementary Figure 3: Comparison of representative 2D class averages from the EMPIAR-10081 dataset obtained using CryoSPARC, RELION and AlignPCA-2D.

### Supplementary Figure 4

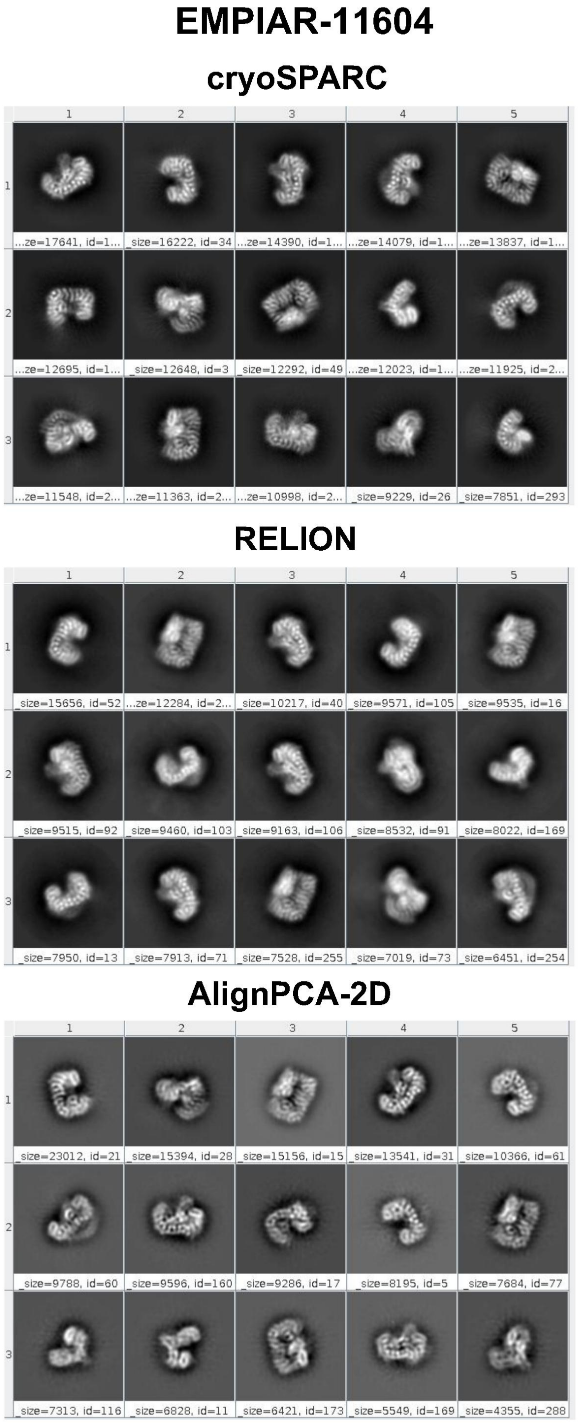


Supplementary Figure 4: Comparison of representative 2D class averages from the EMPIAR-11604 dataset obtained using CryoSPARC, RELION and AlignPCA-2D.

### Supplementary Figure 5

**EMPIAR-10061**

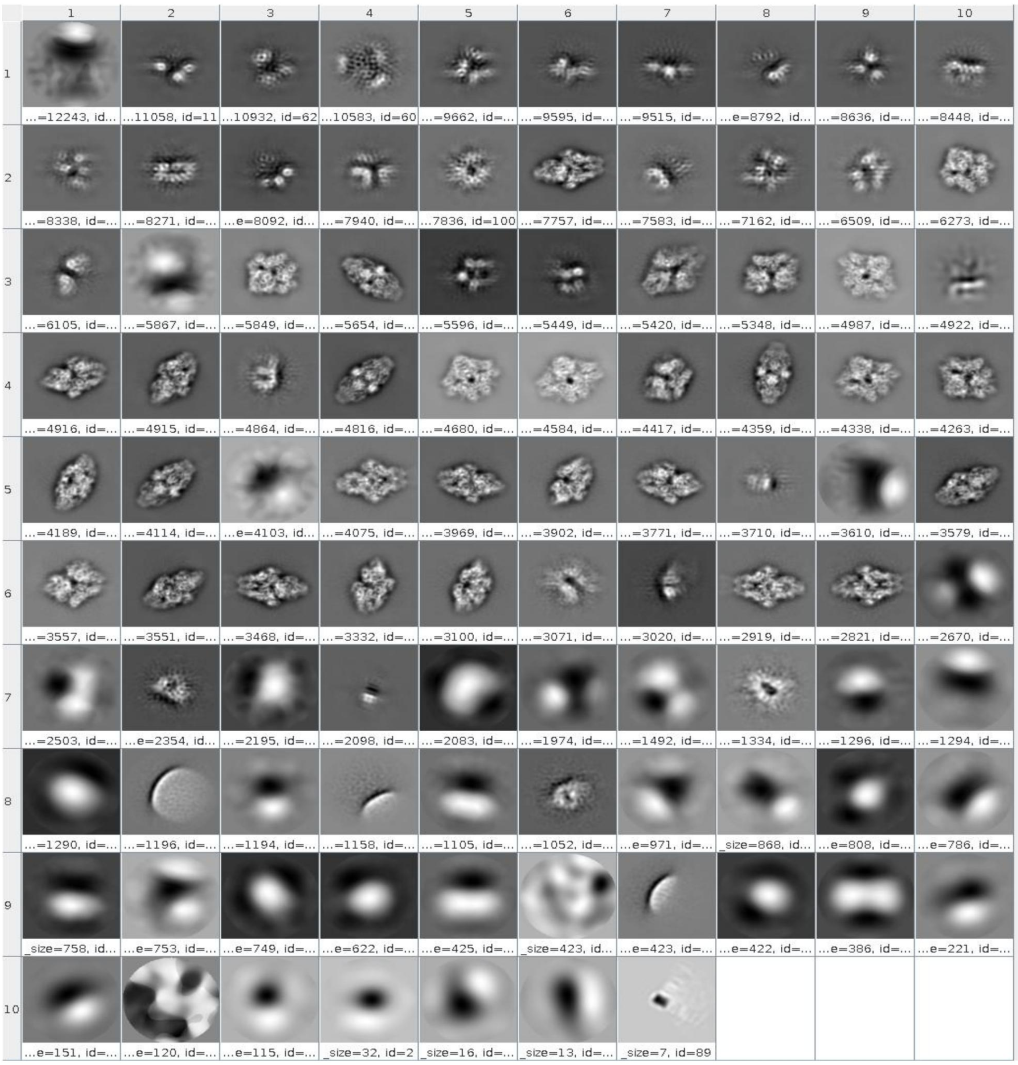


Supplementary Figure 5: Final 2D class averages from the EMPIAR-10061 dataset obtained using AlignPCA-2D.

### Supplementary Figure 6

#### EMPIAR-10025

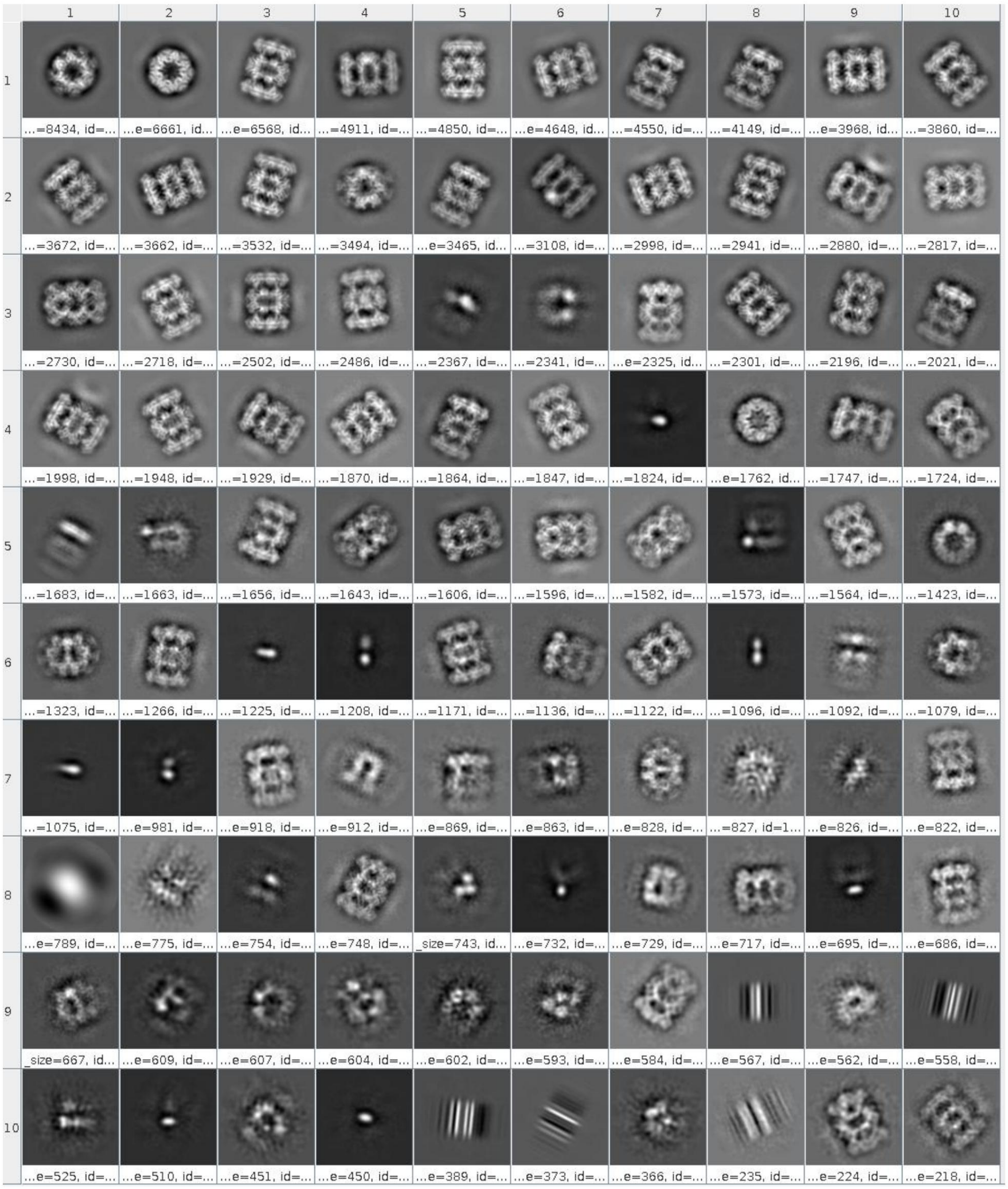


Supplementary Figure 6: Final 2D class averages from the EMPIAR-10025 dataset obtained using AlignPCA-2D.

### Supplementary Figure 7

#### EMPIAR-10081

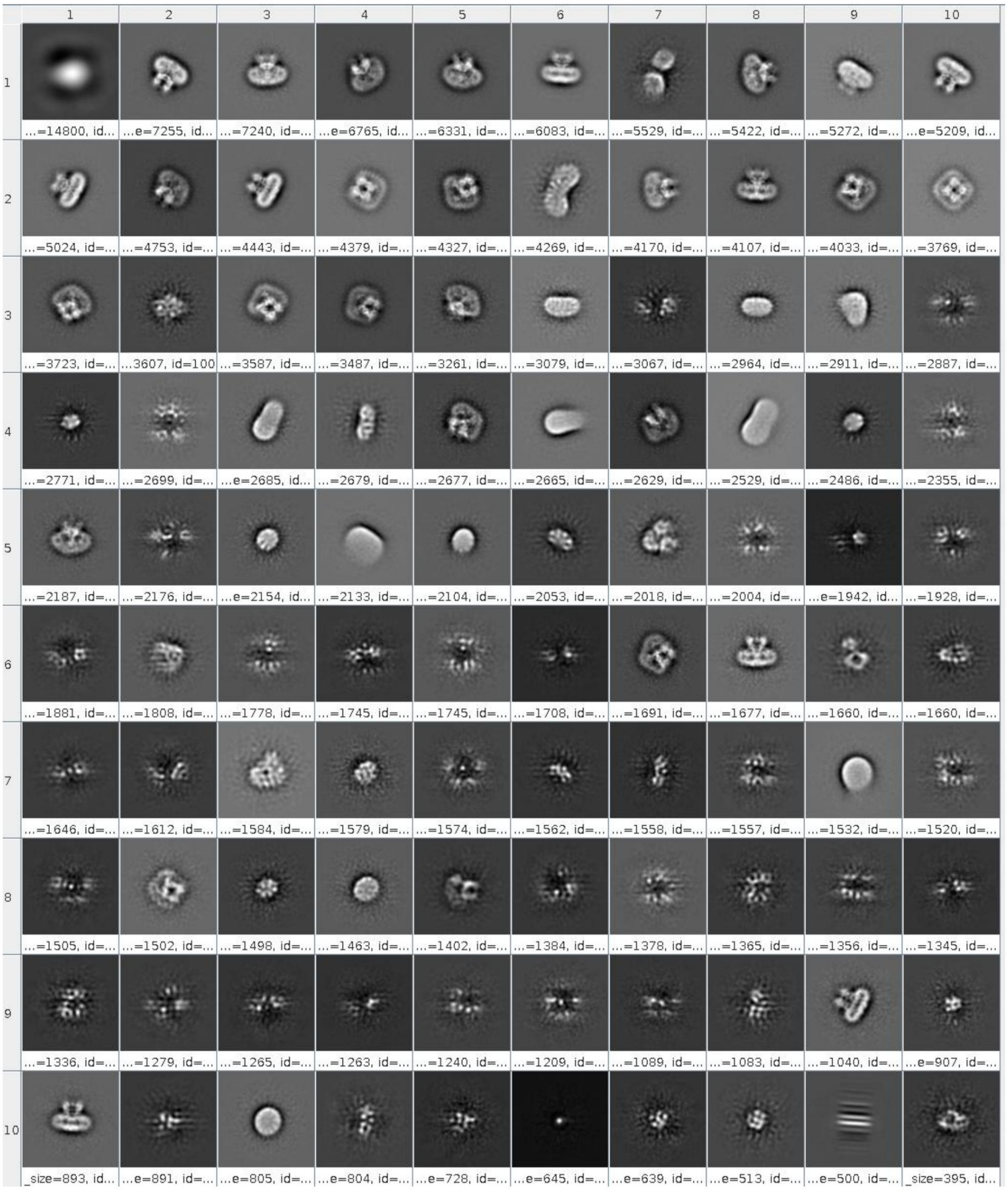


Supplementary Figure 7: Final 2D class averages from the EMPIAR-10081 dataset obtained using AlignPCA-2D.

### Supplementary Figure 8

#### EMPIAR-10061 - CryoSPARC

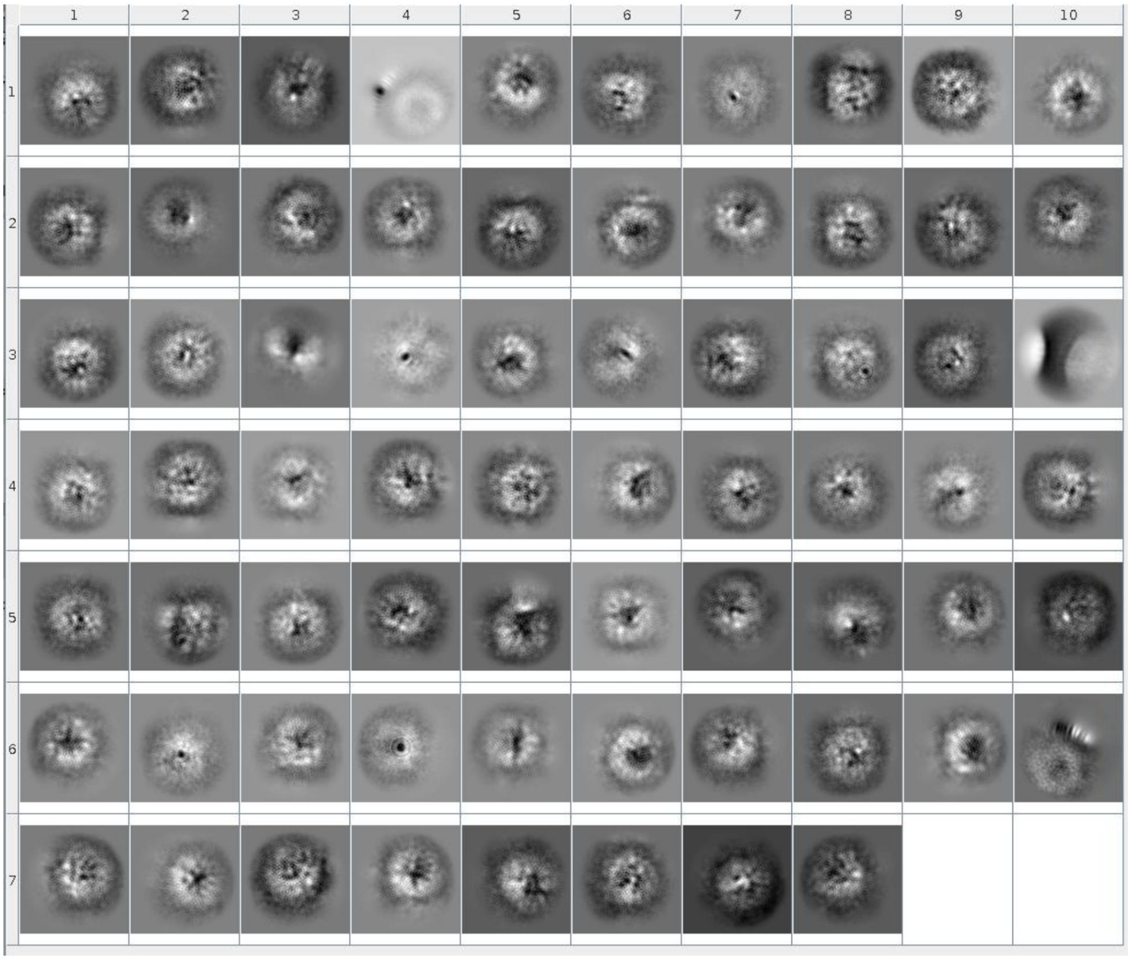


Supplementary Figure 8. Characterization of discarded particles in CryoSPARC. A total of 68 poorly defined classes were identified in the EMPIAR-10061 dataset, representing 212,838 particles.

### Supplementary Figure 9

#### EMPIAR-10061 - rerun CryoSPARC

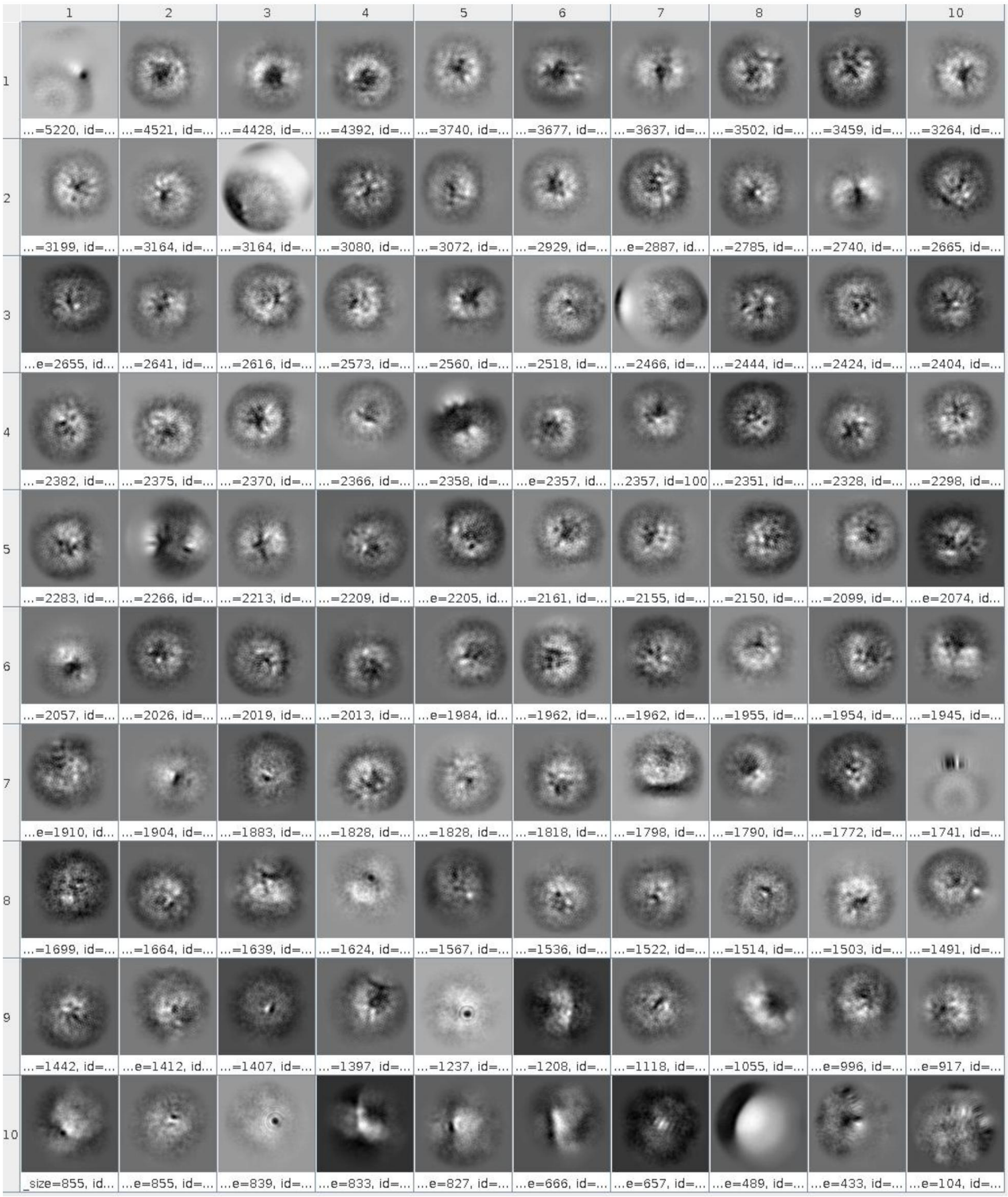


Supplementary Figure 9: Second round of 2D classification in CryoSPARC for dataset EMPIAR-10061. This figure displays the class averages obtained after reprocessing the subset of particles previously identified as poorly defined in Supplementary Figure 8.

### Supplementary Figure 10

#### EMPIAR-10061 - RELION

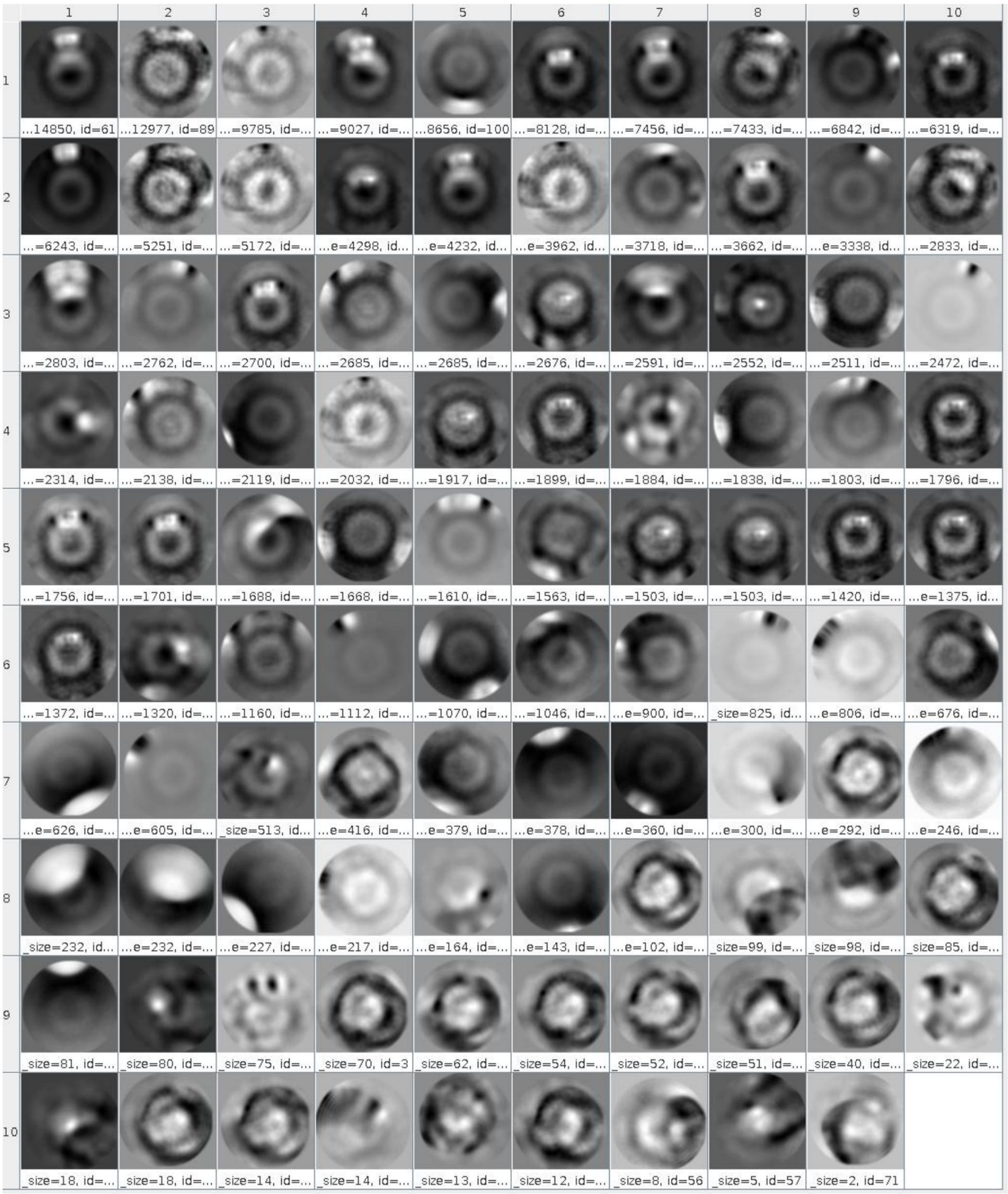


Supplementary Figure 10: Second round of 2D classification with RELION for the EMPIAR-10061 dataset. This figure displays the class averages obtained after reprocessing the subset of particles previously identified as poorly defined by CryoSPARC (as shown in Supplementary Figure 8).

### Supplementary Figure 11

#### EMPIAR-10061 - alignPCA-2D

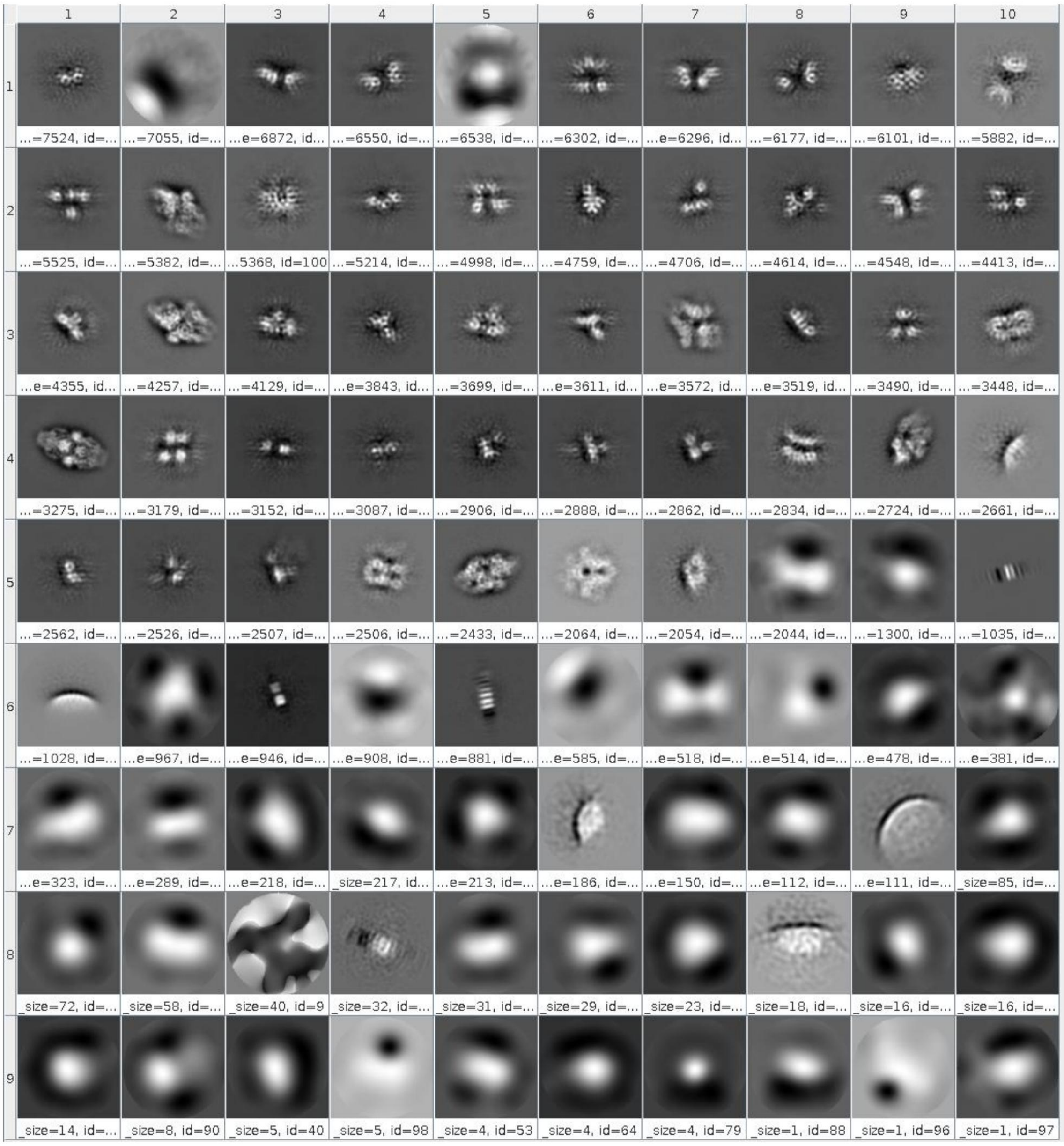


Supplementary Figure 11: Second round of 2D classification with AlignPCA-2D for the EMPIAR-10061 dataset. This figure displays the class averages obtained after reprocessing the subset of particles previously identified as poorly defined by CryoSPARC (as shown in Supplementary Figure 8).

### Supplementary Figure 12

#### EMPIAR-10061 - Align - CryoSPARC

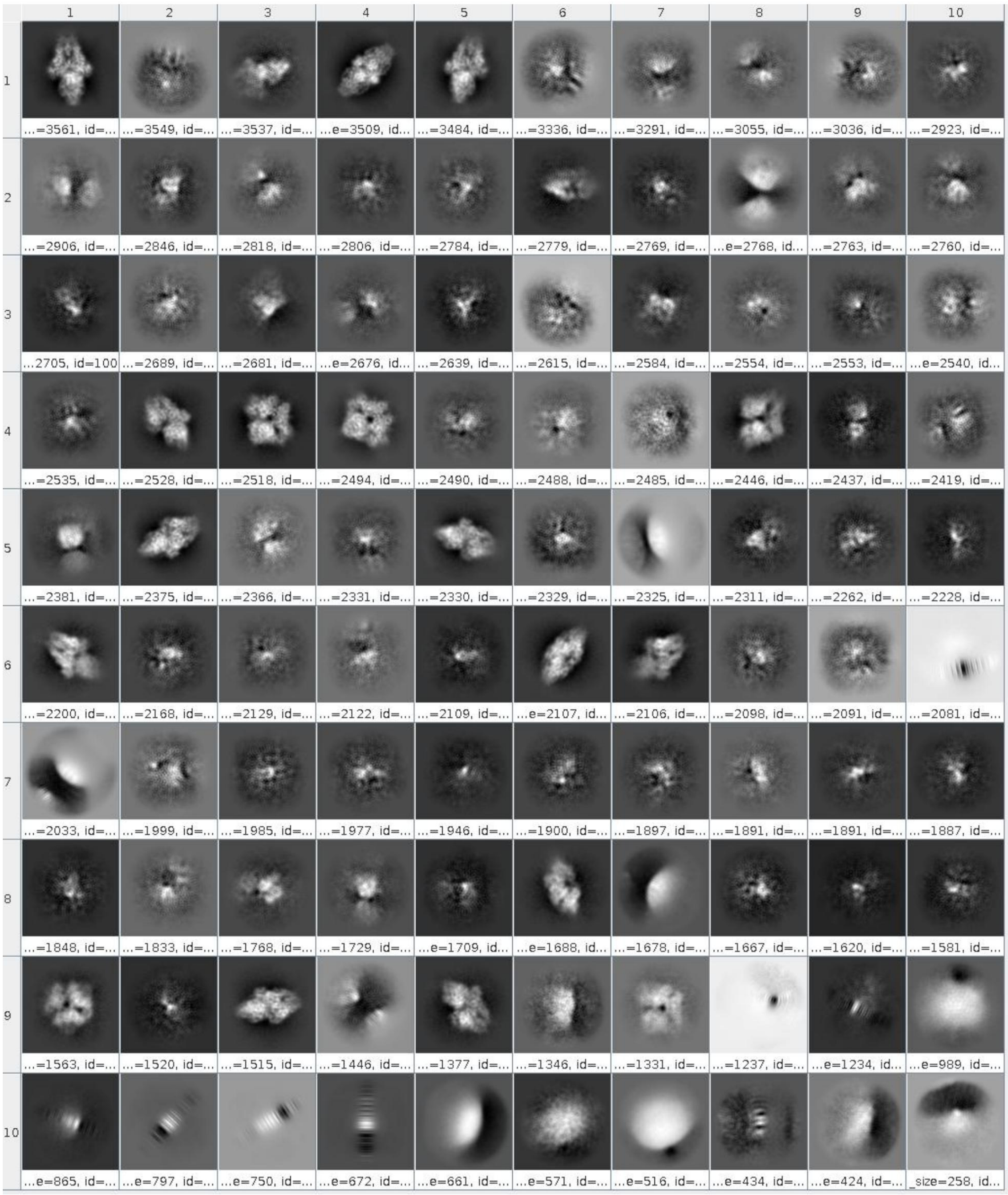


Supplementary Figure 12: Recovery of structural features with CryoSPARC using AlignPCA-2D alignments. The 212,838 particles from the poor-quality classes in Supplementary Figure 8 were re-processed in CryoSPARC. By applying the orientations previously determined by AlignPCA-2D, new 2D class averages were generated, showing improved structural features.

### Supplementary Figure 13

**EMPIAR-10061 - Align - RELION**

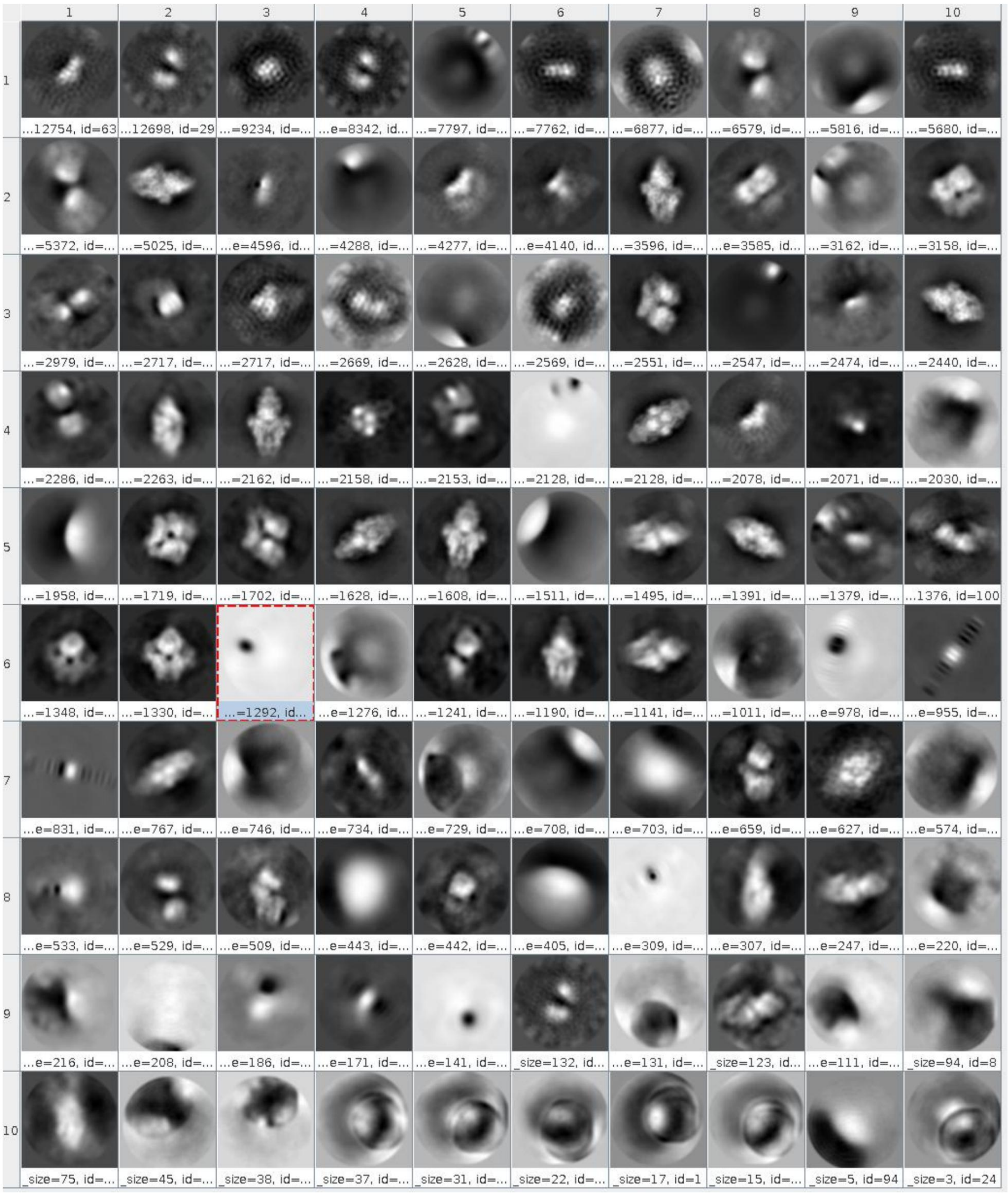


Supplementary Figure 13: Recovery of structural features with RELION using AlignPCA-2D alignments. The 212,838 particles from the poor-quality classes in Supplementary Figure 8 were re-processed in RELION. By applying the orientations previously determined by AlignPCA-2D, new 2D class averages were generated, showing improved structural features.
